# Horizontal gene transfer, segregation loss and the speed of microbial adaptation

**DOI:** 10.1101/2025.03.18.643975

**Authors:** David V. McLeod, Sylvain Gandon

## Abstract

Microbial adaptation is driven by the circulation of mobile genetic elements (MGEs) among bacteria. On the one hand, MGEs can be viewed as selfish genes that spread like infectious diseases in a host population. On the other hand, the horizontal transfer and the loss of these MGEs is often viewed as a form of sexual reproduction that reshuffles genetic diversity in a way that may sometimes be adaptive for bacteria cells. Here, we show how these two perspectives can be reconciled using a single unified framework capturing the dynamics of multiple, interacting MGEs. We apply this framework to study how interactions between MGEs affecting rates of horizontal gene transfer and segregation loss shape the short- and long-term evolutionary dynamics of MGEs and the bacteria population. We show these interactions produce non-random MGE associations that can speed up or slow down microbial adaptation depending on the evolutionary conflicts between MGEs as well as between MGEs and their bacterial hosts. Moreover, we show how these interactions affect the evolutionary potential of the bacteria population. We discuss the implications of these predictions for the community response to environmental stressors such as antibiotic treatment or vaccination campaigns as well as the evolution of accessory genomes.

## 1 Introduction

The horizontal and imperfect vertical transmission of mobile genetic elements (MGEs) plays an important role in microbial adaptation, speeding up or slowing down the spread of MGEs, but also reshuffling genetic material between cells, creating and maintaining cellular diversity. As many of the genes carried by MGEs are important for bacteria survival in variable and challenging environments (e.g., anti-phage defense systems [1, 2], heavy metal resistance [3], antibiotic resistance [3–6]), this cellular diversity likely permits bacteria populations to respond to varied threats [1].

There has been recent interest in understanding how horizontal gene transfer (HGT) and segregation loss (SL) affect the dynamics of microbial adaptation [5, 7–10], and what the implications are for the resilience of bacteria populations [5]. HGT is expected to speed up the evolution of MGE-linked genes by allowing the persistence of costly MGEs [5, 7– 9] and facilitating their rapid dissemination throughout the microbial population [5]. SL, on the other hand, will counteract these processes, slowing the evolution of MGE-linked genes. However, these predictions do not account for how interactions between MGEs affect the rates of HGT and SL [e.g., 11–20]. For example, many MGEs encode defense systems, reducing HGT rates of other MGEs [18], while phage infection can increase HGT rates of integrative conjugative elements [18, 21]. Indeed, in the extreme, some genetic elements cannot transfer autonomously and their horizontal transmission is conditional on the presence of helper MGEs in the same host cell [22]. Such MGE-MGE interactions have the potential to transiently build-up non-random associations between MGEs (linkage disequilibrium [23]), with significant, but poorly understood consequences for microbial adaptation.

Here, we develop a modeling framework to disentangle how MGE-MGE interactions influence HGT and SL and affect microbial adaptation. We show that if MGE-MGE interactions are competitive (interactions reduce HGT and increase SL), MGEs are more likely to be found in different cells, whereas if interactions are mutualistic (increase HGT and reduce SL) and/or MGEs are simultaneously gained and lost, MGEs are more likely to be found in the same cell. In the short-term, such MGE-MGE interactions mean that HGT and SL can speed up or slow down adaptation depending on the evolutionary conflict between MGEs, as well as the conflict between MGEs and bacteria [3, 18, 22, 24]. In the long-term, MGE-MGE interactions affect the persistence of MGEs and the equilibrium distribution of MGE associations. These MGE-MGE associations can have important implications on the ability of the bacteria population to respond to environmental change [25]. We conclude by discussing the implications of our framework for related work concerning the response of bacterial communities to antibiotic treatment [5], the consequences of serotype-specific vaccination [26, 27] and the evolution of accessory genomes [20, 26, 28].

## 2 Model

We first describe our population dynamical model; a schematic of the model is provided in Figure 1 and all notation used is provided in Table 1. Consider a population of bacteria of density *n*(*t*) at time *t*. Bacteria in this population divide at a per-capita rate *b*(1−*n*(*t*)), where *b* is the exponential birth rate, and die at a per-capita rate *d*. Circulating in the bacteria population are two mobile genetic elements (MGEs). Each MGE carries a different gene, denoted *A* and *B*, affecting different aspects of the life-history of the bacteria (e.g., antibiotic resistance [3], virulence factors [29], toxin-antitoxin systems [30], metabolic pathways [31]). Infection by, or carriage of, MGE *X* ∈{*A, B*} (here, and in what follows we identify MGEs by the gene they carry) additively effects the per-capita rate of cell division and bacteria cell death by a total amount *s*_*X*_(*t*). If *s*_*X*_(*t*) *>* 0, gene *X* increases bacterial per-capita growth rate and so carriage of MGE *X* is beneficial. If *s*_*X*_(*t*) *<* 0, gene *X* decreases bacterial per-capita growth rate, and so MGE *X* is parasitic (Box. 1). Each of the *s*_*X*_(*t*) may depend upon population density if, for example, gene *X* affects the rate of cell division, and may also fluctuate in time (e.g., gene *X* controls antibiotic resistance, and antibiotic concentrations temporally vary). We assume genes *A* and *B* do not produce epistasis affecting either the per-capita rate of cell division or cell death. The dynamics of the density of bacteria cells can be written

**Table 1:** Notation used in main text.

| Symbol | Description |
| --- | --- |
| $n(t)$ | Total density of bacteria cells. |
| $b(1 - n(t)), d$ | Per-capita rate of bacterial cell division and cell death, respectively. |
| $s_X(t)$ | Additive effect carriage of MGE $X \in \{A, B\}$ has on the per-capita rate of cell division and bacteria cell death. |
| $p_X(t), q_X(t)$ | Frequency of cells carrying, and not carrying, respectively, MGE $X$. |
| $p_{AB}(t)$ | Frequency of cells carrying both MGEs. |
| $D(t)$ | Linkage disequilibrium between MGEs. |
| $\tilde{\theta}_X, \theta_X$ | Probability per cell division of SL of only MGE $X$ by a cell carrying, and not carrying, respectively, MGE $Y$. |
| $\theta_{AB}$ | Probability per cell division of SL of both MGEs. |
| $\theta_X^Y \equiv \tilde{\theta}_X + \theta_{AB}$ | Probability per cell division of SL of MGE $X$ by a cell carrying both MGEs. |
| $\langle \theta_X \rangle$ | Probability per cell division of SL of MGE $X$ by a randomly selected cell carrying MGE $X$. |
| $\langle \theta_X \rangle_0$ | $\langle \theta_X \rangle$ evaluated when $D(t) = 0$. |
| $\tilde{\beta}_X, \beta_X$ | Rate of transmission of only MGE $X$ by a donor carrying, and not carrying, respectively, the other MGE $Y$. |
| $\beta_{AB}$ | Rate of simultaneous transmission of both MGEs $A$ and $B$. |
| $\beta_X^Y = \tilde{\beta}_X + \beta_{AB}$ | Rate of transmission of MGE $X$ if the donor carries both MGEs. |
| $\langle \beta_X \rangle$ | Average per-capita rate of transmission of MGE $X$ by a randomly selected cell. |
| $\langle \beta_X \rangle_0$ | $\langle \beta_X \rangle$ evaluated when $D(t) = 0$. |
| $\lambda_X^Y, \lambda_X$ | Probability MGE $X$ successfully establishes in a recipient carrying, and not carrying, respectively, the other MGE $Y$. |
| $\langle \lambda_X \rangle$ | Probability MGE $X$ successfully establishes in a randomly selected host that does not carry MGE $X$. |
| $\langle \lambda_X \rangle_0$ | $\langle \lambda_X \rangle$ evaluated when $D(t) = 0$. |

**Figure 1:**
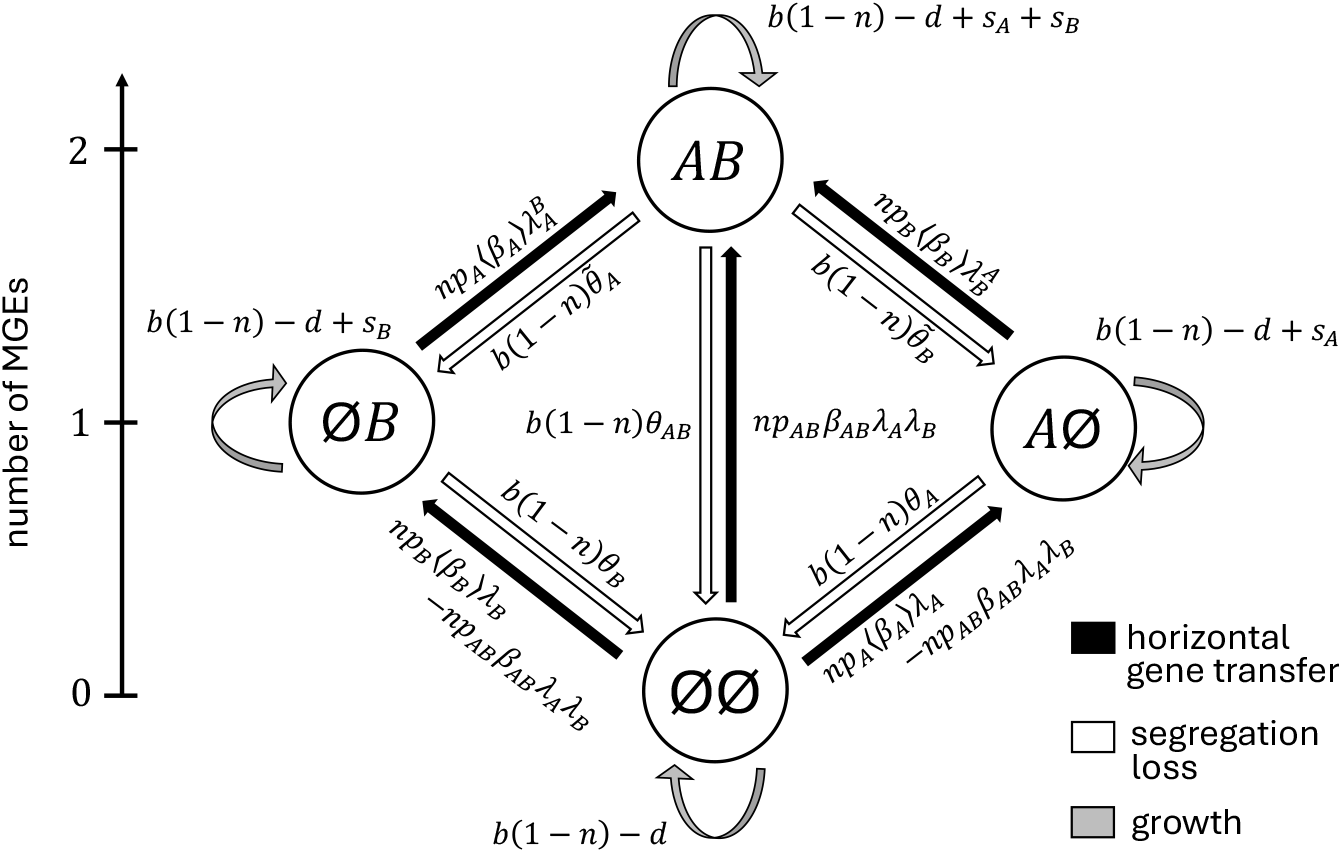
Schematic of population dynamics. We consider a population of bacteria cells and two MGEs carrying different genes, denoted *A* and *B*, affecting the bacteria life cycle. There are four bacterial states, or genotypes, *ij* ∈ {∅∅ , *A*∅ , ∅*B* , *AB*}, where ∅ indicates the absence of the MGE. We let *p*_*X*_(*t*) denote the frequency of MGE *X*, and *p*_*AB*_(*t*) denote the frequency of cells carrying both MGEs. The density of bacteria cells depends on the per-capita growth rates, whereas the distribution of MGEs in the population depends on the per-capita growth rates as well as HGT and SL. All notation used is defined in Table 1.

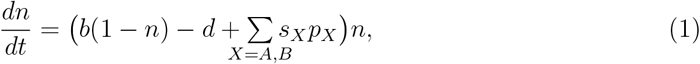

where *p*_*X*_(*t*) is the frequency of bacteria cells carrying MGE *X* at time *t*.

In addition to affecting the per-capita rates of cell division and bacteria cell death, MGEs may be shuffled between cells and lost. First, MGEs may fail to be inherited during cell division due to segregation loss (SL). SL of an MGE may be affected by the presence of the other MGE [32] (e.g., due to plasmid incompatibilities [33–35]), as well as whether multiple MGEs can be lost during the same cell division. Therefore, let *θ*_*X*_ denote the probability MGE *X* is lost from a cell that does not carry the other MGE *Y* , and 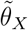 the probability that *only* MGE *X* is lost from a cell that does carry the other MGE *Y* . Let *θ*_*AB*_ denote the probability that a cell loses both MGEs during the same cell division. Thus, the probability that MGE *X* is lost from a cell carrying both MGEs is 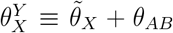. Consequently, if 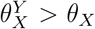 (resp. 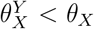), MGE *X* is more (resp. less) likely to be lost if the dividing cell also carries the other MGE. The probability a cell carrying both MGEs loses *at least* one is 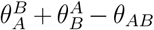. More generally, the probability of SL per cell division of a randomly selected cell carrying MGE *A* is

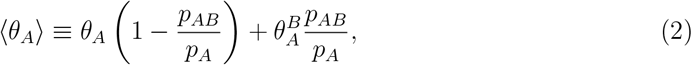

where *p*_*AB*_ is the frequency of cells carrying both MGEs. A similar expression can be derived for MGE *B*.

Second, MGEs may be transmitted from one cell (the “donor”) to another cell (the “recipient”) through horizontal gene transfer (HGT). As the presence of other MGEs in the donor and/or recipient can both facilitate and inhibit rates of HGT [11–18] (see also [19] for a comprehensive review for plasmids), we decompose HGT into two parts: (1) the transmission of the MGE(s) from the donor, and (2) the successful establishment of the transmitted MGE(s) in the recipient. Transmission of the MGE(s) captures the rate of production, or mobility, of the MGE. Some of this mobility is governed by extrinsic factors, such as the presence of phage for transduction, but it may also be driven by intrinsic factors, such as the machinery needed for conjugation which can depend on the plasmid as well as the host. Thus the transmission of the MGE(s) may be affected by whether the donor carries multiple MGEs, and if so, whether one or both are transmitted (e.g., co-transfer of plasmids [14]). Let 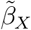 and *β*_*X*_ denote the transmission rate of only MGE *X* if the donor does, and does not, respectively, carry the other MGE *Y* . Let *β*_*AB*_ denote the rate of transmission of both MGEs *A* and *B*. If the donor carries both MGEs, the rate of transmission of MGE *X* is 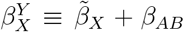, while the rate at which such a cell transmits *at least* one MGE is 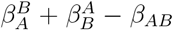. Hence, if 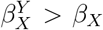 (resp. 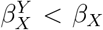), MGE *X* is more (resp. less) likely to be transmitted if the donor carries the other MGE. More generally, the average rate of transmission of MGE *A* by a randomly selected cell carrying MGE *A* is *n*⟨*β*_*A*_⟩, where

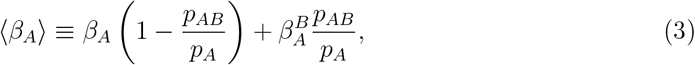

and similarly for MGE *B*.

Successful establishment of the transmitted MGE(s) may depend on the genes carried by the MGEs present in the recipient, for example, MGE-linked defense systems [18]. Let 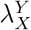 denote the probability MGE *X* successfully establishes if the recipient carries MGE *Y* , and let *λ*_*X*_ denote the probability MGE *X* establishes otherwise. We assume the probabilities of establishment are independent. Hence, if two MGEs are acquired at the same time their probability of being maintained in the lineage is independent of the other. Thus the probability both MGEs are established in an *ab* host, given they were transmitted, is *λ*_*A*_*λ*_*B*_. Finally, we ignore “multicopy” effects, and so if a MGE is transmitted to a host who already carries that MGE, the state of the recipient host is unchanged, irrespective of whether establishment occurs or not. The probability of establishment of MGE *A* in a randomly selected host that does not carry MGE *A* is

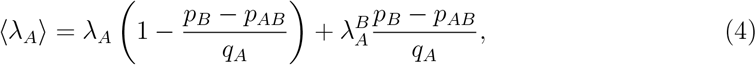

where *q*_*A*_(*t*) = 1 − *p*_*A*_(*t*), and similarly for MGE *B*. Under these assumptions, the population dynamics are presented in Figure 1, while the equations describing the dynamics of the densities of each type of bacteria cell can be found in Appendix A.

As SL and HGT do not affect the rate of cell division or bacteria cell death, they will only affect the dynamics of the total density of bacteria cells via their effects on the change in frequency *p*_*X*_(*t*) of the different MGEs (equation (1)). However, SL and HGT can affect how MGEs are distributed between cells by creating nonrandom associations between MGEs. Such nonrandom associations can be, quantified by the variable *D*(*t*) which measures the linkage disequilibrium (LD) between MGEs:

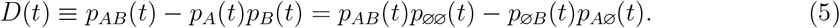

Here, *p*_*ij*_(*t*) is the frequency of cells with the set of MGE(s) *ij* ∈ {∅∅ , *A*∅ , ∅*B* , *AB*} , where ∅ indicates the absence of the MGE. The linkage disequilibrium between MGEs has a straightforward interpretation: if *D*(*t*) *>* 0, cells are more likely to carry either both MGEs or neither, whereas if *D*(*t*) *<* 0, cells are more likely to carry one MGE. Note that each of the *p*_*ij*_(*t*) can be expressed using solely the variables for MGE frequencies and *D*(*t*).

## 3 Results

Evolution of gene *X* corresponds to a change in its frequency, captured by the differential equation

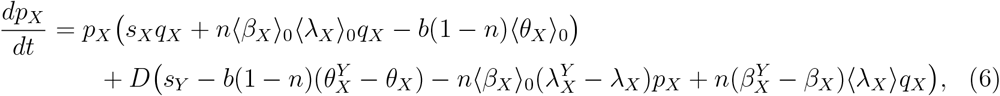

where we use the notation ⟨*Z*_*X*_⟩_0_ for *Z* ∈ {*β, λ, θ*} to indicate we have set *D*(*t*) = 0 in the quantity ⟨*Z*_*X*_⟩ using equations (2)-(4). Thus the quantities ⟨*Z*_*X*_⟩ _0_ depend solely on the frequency of the other MGE *Y* . For example, if *Z* = *β* and *X* = *A*, then setting *D*(*t*) = 0 in equation (3) yields

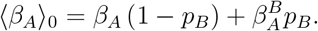

This notation allows us to highlight that the evolutionary dynamics of gene *X* in equation (6) has two components, which we will refer to as the *direct effect* and the *indirect effect*. The direct effect captures how gene *X* directly alters its evolution without accounting for any non-random associations with the other MGE-linked gene. It is the first line on the right-hand side of (6), and it consists of three factors. The first factor, *s*_*X*_(*t*)*p*_*X*_(*t*)*q*_*X*_(*t*), represents direct selection on gene *X*: if *s*_*X*_(*t*) *>* 0, gene *X* is advantageous and so direct selection on gene *X* will increase its frequency. The second factor, *n*⟨*β*_*X*_⟩ _0_ ⟨*λ*_*X*_⟩ _0_*p*_*X*_(*t*)*q*_*X*_(*t*), represents how HGT directly increases the frequency of gene *X* when it is carried by an MGE. Finally, the third factor, −*b*(1 − *n*) ⟨*θ*_*X*_⟩ _0_*p*_*X*_(*t*), represents how SL directly decreases the frequency of gene *X* when it is carried by an MGE. The indirect effect captures how non-random associations between gene *X* and the other gene *Y* affect the dynamics of gene *X*. It is the second line on the right-hand side of (6).

Before considering the indirect effect in detail, note that if MGE *X* has decreased SL 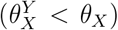, increased transmission 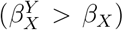 and/or increased establishment probability 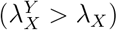 in the presence of the other MGE *Y* , then MGE *Y* increases the fitness of MGE *X* through HGT and SL. We will therefore refer to this as *mutualistic HGT and SL*, or a *mutualism* between MGEs (Box. 1). If instead MGE *X* has increased SL 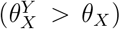, decreased transmission 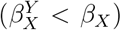 and/or decreased establishment probability 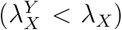 in the presence of MGE *Y* , MGE *Y* decreases the fitness of MGE *X* through HGT and SL. We will therefore refer to this as *competitive HGT and SL* or *competition* between MGEs (Box. 1). While it is biologically possible that HGT and SL show a mixture of competitive and mutualistic effects, for simplicity our analysis will focus upon situations in which they are either all competitive (or neutral), or all mutualistic (or neutral).

### 3.1 Generation of the indirect effect through HGT and SL

In a large, unstructured population with no epistatic interactions between genes *A* and *B*, the expectation is that the genes *A* and *B* will be in linkage equilibrium (i.e., *D*(*t*) = 0; [36–38]), and so the indirect effect will be zero. Indeed, if we calculate the dynamical equation for *D*(*t*) in the absence of MGE-MGE interactions we obtain

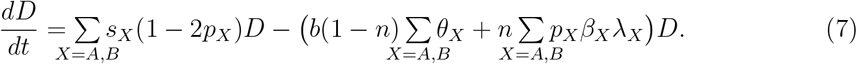

Thus in the absence of MGE-MGE interactions, HGT and SL act to remove *D*(*t*) from the population, similar to genetic recombination and/or mutation. Therefore, we ask when MGE-MGE interactions generate *D*(*t*). To answer this, we derive the dynamical equation for *D*(*t*) in the presence of MGE-MGE interactions (see Appendix A.1) and then we set *D*(*t*) = 0, giving

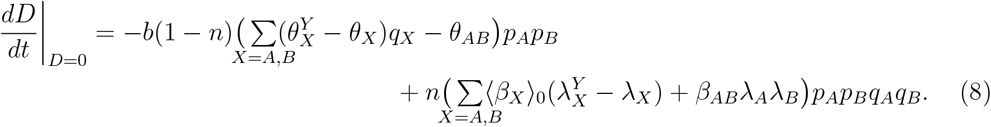

Thus MGE-MGE interactions affecting HGT and SL can generate both an over-representation of cells carrying both MGEs or neither (*D*(*t*) *>* 0) or an over-representation of cells carrying a single MGE (*D*(*t*) *<* 0). We provide a geometric explanation of the logic in Figure 2; here we focus upon the key predictions.

**Figure 2:**
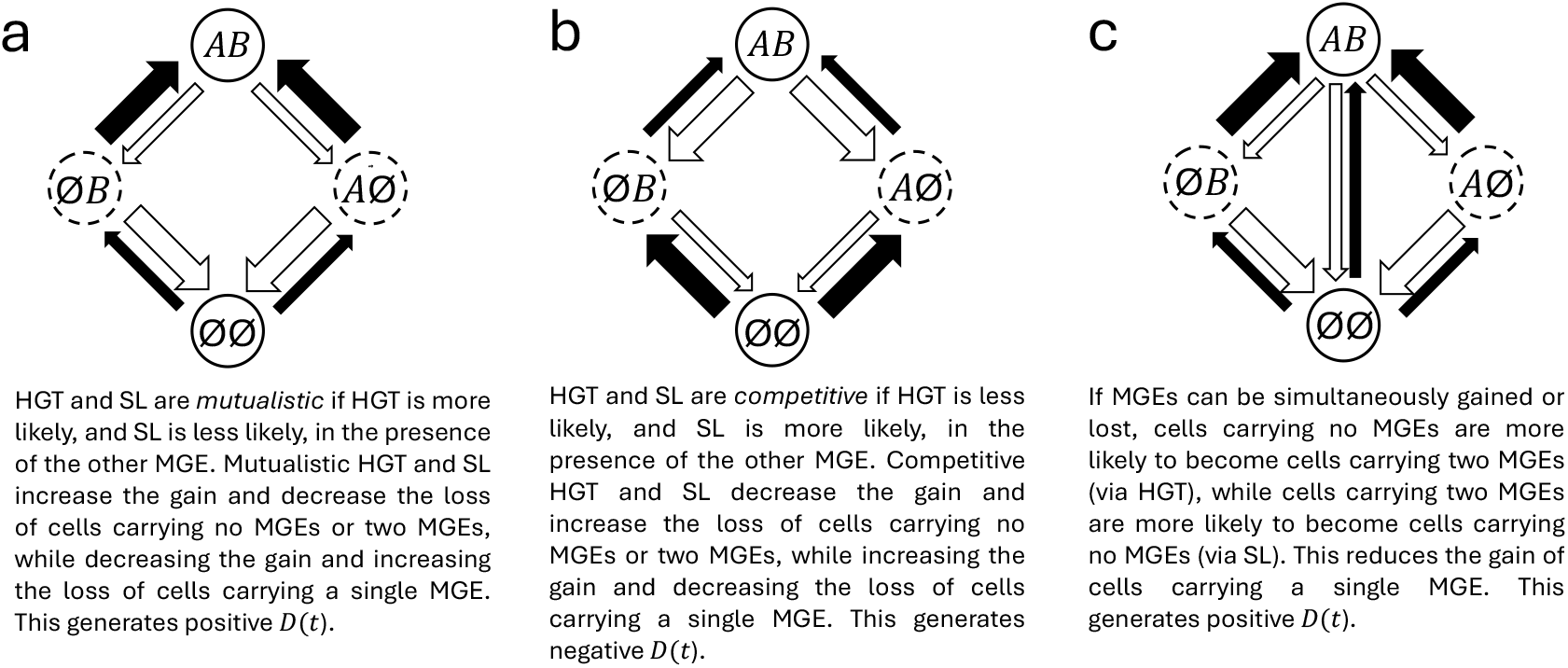
MGE-MGE interactions affecting HGT and SL generate. *D*(*t*). Illustration of the logic behind how MGE-MGE interactions shape the distribution of MGEs within the bacteria population, *D*(*t*). Since by definition *D*(*t*) = *p*_*AB*_(*t*)*p*_∅∅_(*t*)−*p*_*A*∅_(*t*)*p*_∅*B*_(*t*), any process that increases the frequency of ∅∅ and *AB* cells (solid circles) will generate positive *D*(*t*), whereas any process that increases the frequency of ∅*B* and *A*∅ cells (dashed circles) will generate negative *D*(*t*).

#### Competitive HGT and SL tends to generate *D*(*t*) *<* 0 while mutualistic HGT/SL generates *D*(*t*) *>* 0

From inspection of (8), mutualistic HGT and SL will tend to produce an over-representation of cells carrying both MGEs or neither (positive *D*(*t*)), whereas competitive HGT and SL will tend to produce an over-representation of cells carrying one MGE (negative *D*(*t*)). Because mutualistic or competitive transmission 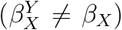 simply affects the increase in frequency of MGEs and not the identity of the recipient cells, the recipient is likely to carry MGE *Y* with a probability equal to its existing frequency, *p*_*Y*_ (*t*). Consequently, mutualistic or competitive transmission on its own cannot generate nonrandom associations between MGEs (i.e., nonzero *D*(*t*)).

#### The simultaneous loss and gain of MGEs produces positive *D*(*t*)

If MGEs can be simultaneously lost (*θ*_*AB*_ *>* 0) and/or gained (*β*_*AB*_), from (8) this will generate positive *D*(*t*). Indeed, from equation (8), SL will produce positive *D*(*t*) if

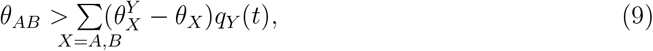

whereas HGT will produce positive *D*(*t*) if

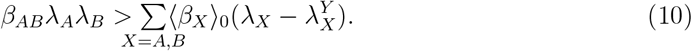

Thus, the simultaneous gain and loss increases the likelihood of positive *D*(*t*), independent of whether HGT and SL are competitive or mutualistic. However, from (9) and (10), if HGT and SL are competitive or mutualistic and the simultaneous loss and gain of MGEs is possible, MGE frequency becomes increasingly important. For example, from (9), as MGEs become increasingly frequent (*q*_*Y*_ (*t*) → 0), whether SL is competitive or mutualistic matters less, and positive *D*(*t*) is always produced, while as MGEs become increasingly rare (*q*_*Y*_ (*t*) → 1), such interactions are increasingly important (Appendix A.2).

### 3.2 The indirect effect and the speed of microbial adaptation

Next we ask how the indirect effects generated by the MGE-MGE interactions alter microbial adaptation via the change in frequency of MGE *X*. To answer this, note that the indirect effect of (6) consists of two components. The first is the presence of terms affecting rates of HGT and SL, while the second is indirect selection on MGE-linked gene *Y* .

### The indirect effect of competitive or mutualistic SL reduces the loss of MGEs

SL always leads to a decrease in the frequency of gene *X*, that is, the combined direct and indirect effect of SL always reduces MGE frequency. What is less clear is how MGE-MGE SL interactions, and their generation of *D*(*t*), affect the speed of evolution of gene *X* as compared to the situation if *D*(*t*) is identically zero. That is, we are comparing two models which have the same direct effect, but only one has the indirect effect. If SL is competitive or mutualistic, from (6) the indirect effect of SL will increase the frequency of MGE *X* if

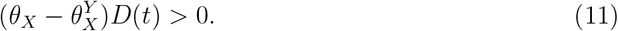

Since 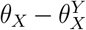 generates same sign *D*(*t*) (Fig. 3**a**,**b**), from inequality (11), the indirect effect of competitive or mutualistic SL will increase the frequency of MGE *X*, and so slow the loss of MGE *X* by SL (Fig. 3**a**,**c**). If MGEs can be simultaneously lost, although this increases the likelihood of generating positive *D*(*t*), as *θ*_*AB*_ does not independently appear in equation (6) it will not otherwise affect the change in frequency of MGE *X* (Fig. 3**a**)

**Figure 3:**
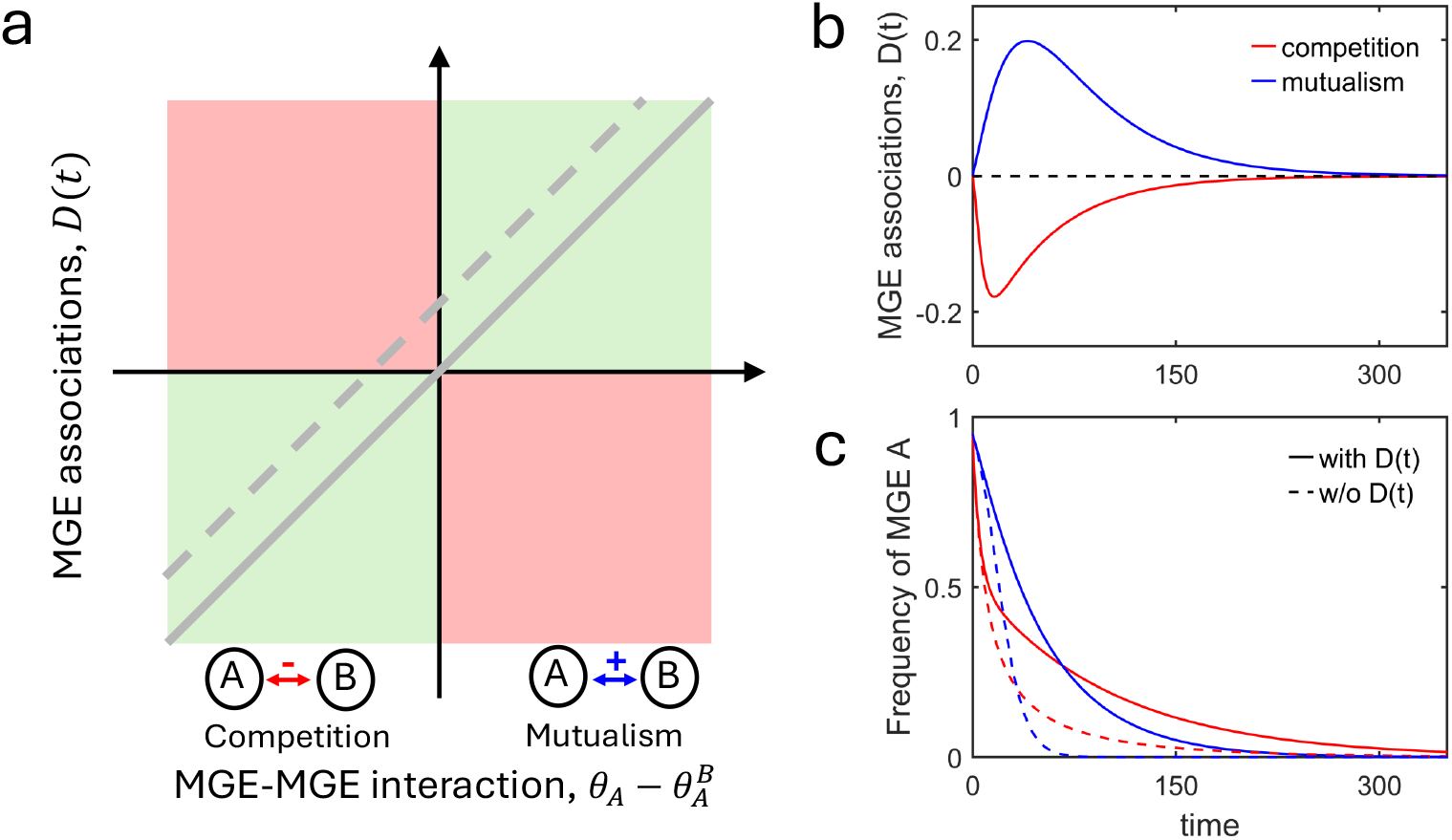
Indirect effect of MGE-MGE interactions on SL (in the absence of HGT) on the change in frequency of MGE. **A**. In the absence of HGT, equation (6) consists of the direct effect plus the term 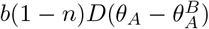. Since MGE-MGE interactions (sign of 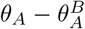) generate same sign *D*(*t*) (grey line in panel **a**; see panel **b**), the system stays in the “green zone”, where the indirect effect increases the frequency of MGE *A* (panel **c**). The simultaneous loss of MGEs generates positive *D*(*t*), independently of MGE-MGE interactions (dashed grey line). If interactions are weakly competitive (dashed line is in red zone; panel **a**) the above predictions may be reversed. In panel **b**,**c**, all simulations assume both genes are neutral, *s*_*X*_ = 0, there is no HGT, 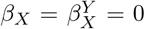, and use parameter values *b* = 4, *d* = 2, *θ*_*AB*_ = 0, with initial conditions *p*_*A*_(0) = 0.95, *p*_*B*_(0) = 0.95, *D*(0) = 0, *n*(0) = (*b*−*d*)*/b*. If SL is competitive (red curves), 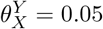 and *θ*_*X*_ = 0.005, whereas if SL is mutualistic (blue curves), 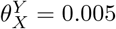 and *θ*_*X*_ = 0.05.

#### The indirect effect of competitive or mutualistic HGT can reduce or enhance the gain of MGEs

HGT always leads to an increase in the frequency of MGE *X*, that is, the combined direct and indirect effect of HGT always increases MGE frequency. To unravel the indirect effect, focus first on those aspects of HGT capable of generating *D*(*t*) (so set 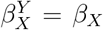). In this case, if HGT is competitive or mutualistic, from (6) the indirect effect of HGT will decrease the frequency of MGE *X* if

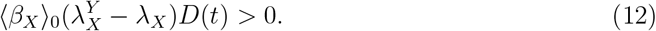

Since 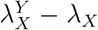 generates same sign *D*(*t*), the indirect effect of competitive or mutualistic HGT will decrease the frequency of MGE *X*, and so slow the evolution of gene *X* by HGT (Fig. 4**a**,**b**). If MGEs can be simultaneously transmitted, although *β*_*AB*_ *>* 0 increases the likelihood of positive *D*(*t*), it does not independently appear in equation (6). Therefore it will not otherwise affect the change in frequency of MGE *X* (Fig. 4**a**)

**Figure 4:**
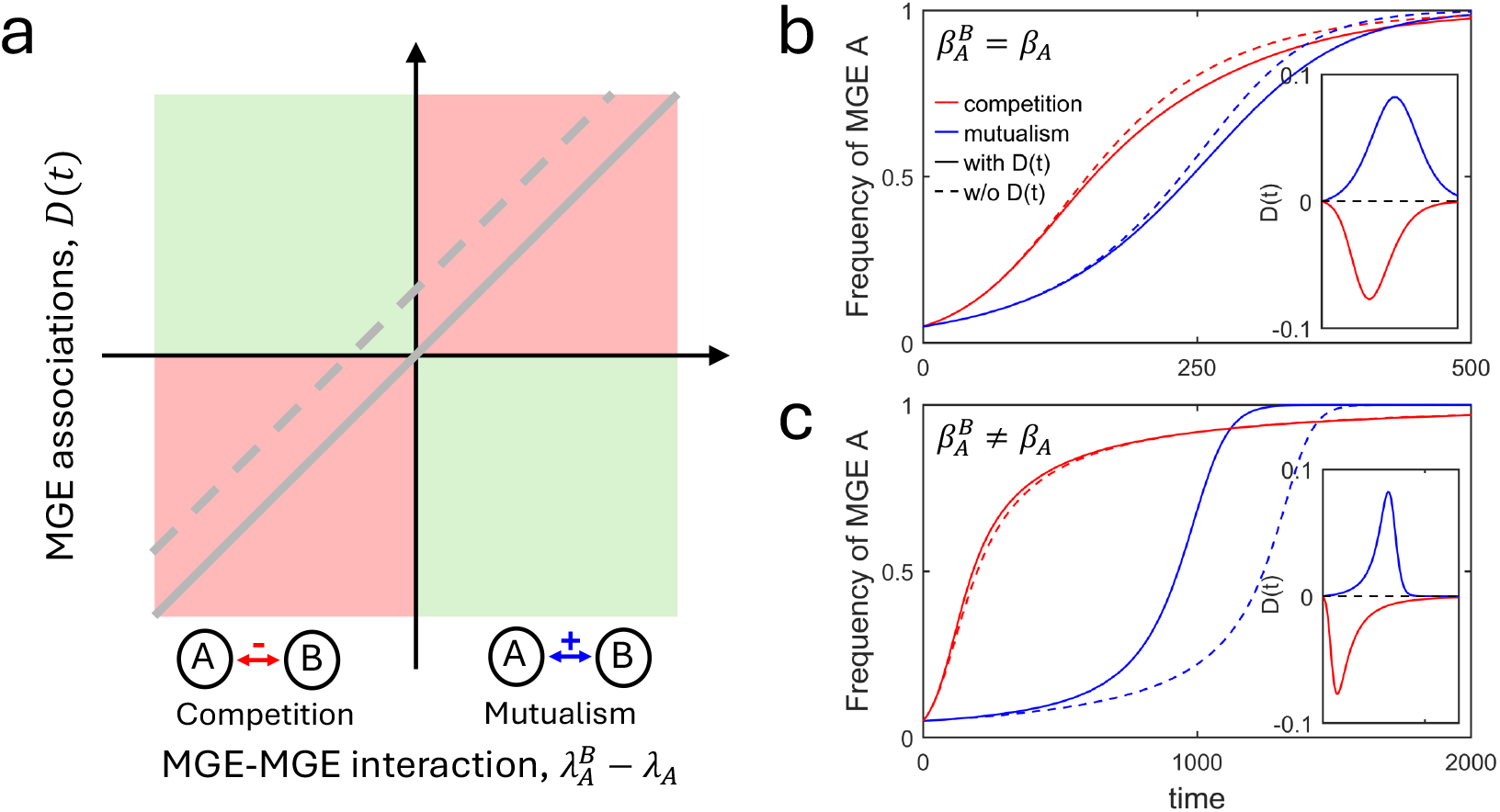
Indirect effect of MGE-MGE interactions on HGT (in the absence of SL) on the change in frequency of MGE. **A.** In the absence of SL, equation (6) consists of the direct effect plus the term –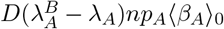 and the term 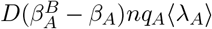. MGE-MGE interactions (sign of 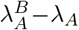) generate same sign *D*(*t*) (grey line, panel **a**), so if 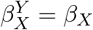, the system stays in the “red zone” where the indirect effect decreases the frequency of MGE *X* (panel **a**,**b**). The simultaneous gain of MGEs generates positive *D*(*t*), independently of MGE-MGE interactions (dashed grey line). If interactions are weakly competitive, the above predictions may be reversed. If transmission dependencies, 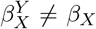, are sufficiently strong and MGE *X* is not too abundant, the red and green zones are reversed, and so the indirect effect increases the frequency of MGE *X* (panel **c**). In panel **b**,**c**, all simulations assume both genes are neutral, s_*X*_ = 0, there is no SL,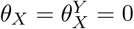, and use parameter values *b* = 4, *d* = 2, *β*_*AB*_ = 0 with initial conditions *p*_*A*_(0) = 0.05, *p*_*B*_(0) = 0.05, *D*(0) = 0, *n*(0) = (*b* − *d*)*/b*. If HGT is competitive (red curves), 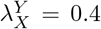 and *λ*_*X*_ = 0.9, whereas if HGT is mutualistic (blue curves), 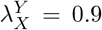 and *λ*_*X*_ = 0.4. In panel **b**, 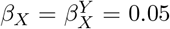, while in panel **c** 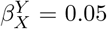 and *β*_*X*_ = 0.0025 (mutualistic HGT) or 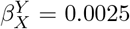 and *β*_*X*_ = 0.05 (competitive HGT).

Competitive or mutualistic HGT need not be restricted to 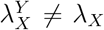, and can also involve interactions affecting transmission, 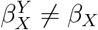. While MGE-MGE interactions affecting transmission cannot generate *D*(*t*), they can affect the speed of evolution of gene *X* once *D*(*t*) has been generated. In particular, the indirect effect of competitive or mutualistic HGT will increase the frequency of MGE *X* if

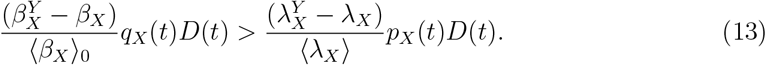

Thus if transmission interactions are sufficiently strong, both competitive and mutualistic HGT cause the indirect effect of HGT to increase the frequency of MGE *X* (Fig. 4**c**).

#### Indirect selection and MGE-MGE and MGE-bacteria conflict

Recall that in addition to affecting the rates of HGT and SL, from equation (6) the indirect effect also involves indirect selection on the other MGE-linked gene *Y* , *s*_*Y*_ *D*(*t*). If *s*_*Y*_ and *D*(*t*) share sign, indirect selection will increase the frequency of MGE *X*, whereas if they are of opposite sign indirect selection will decrease the frequency of MGE *X*. The question, therefore, is how MGE-MGE interactions affecting HGT and SL determine the relationship between the sign of *s*_*Y*_ and *D*(*t*). This can be conceptually thought of as whether there is conflict between the MGEs (i.e., are HGT and SL competitive or mutualistic?), and whether there is conflict between the MGEs and bacteria cells (Fig. 5; [3, 18, 22, 24]). If carriage of MGE *Y* decreases bacteria cell fitness (*s*_*Y*_ *<* 0), bacteria and MGE *Y* are in conflict, whereas if it increases bacterial cell fitness (*s*_*Y*_ *>* 0), they are not. Since mutualistic HGT and SL produces positive *D*(*t*) while competitive HGT and SL produce negative *D*(*t*), indirect selection will increase the frequency of MGE *X* if either there are evolutionary conflicts between all parties (MGE *A, B*, and bacteria cell), or all parties are mutualistic (Fig. 5). If instead conflict occurs at one level (e.g., competition between MGEs) while a mutualistic interaction occurs at the other (e.g., MGE *Y* benefits the bacteria cell), the generation of indirect selection by HGT and SL decreases the frequency of MGE *X* (Fig. 5).

**Figure 5:**
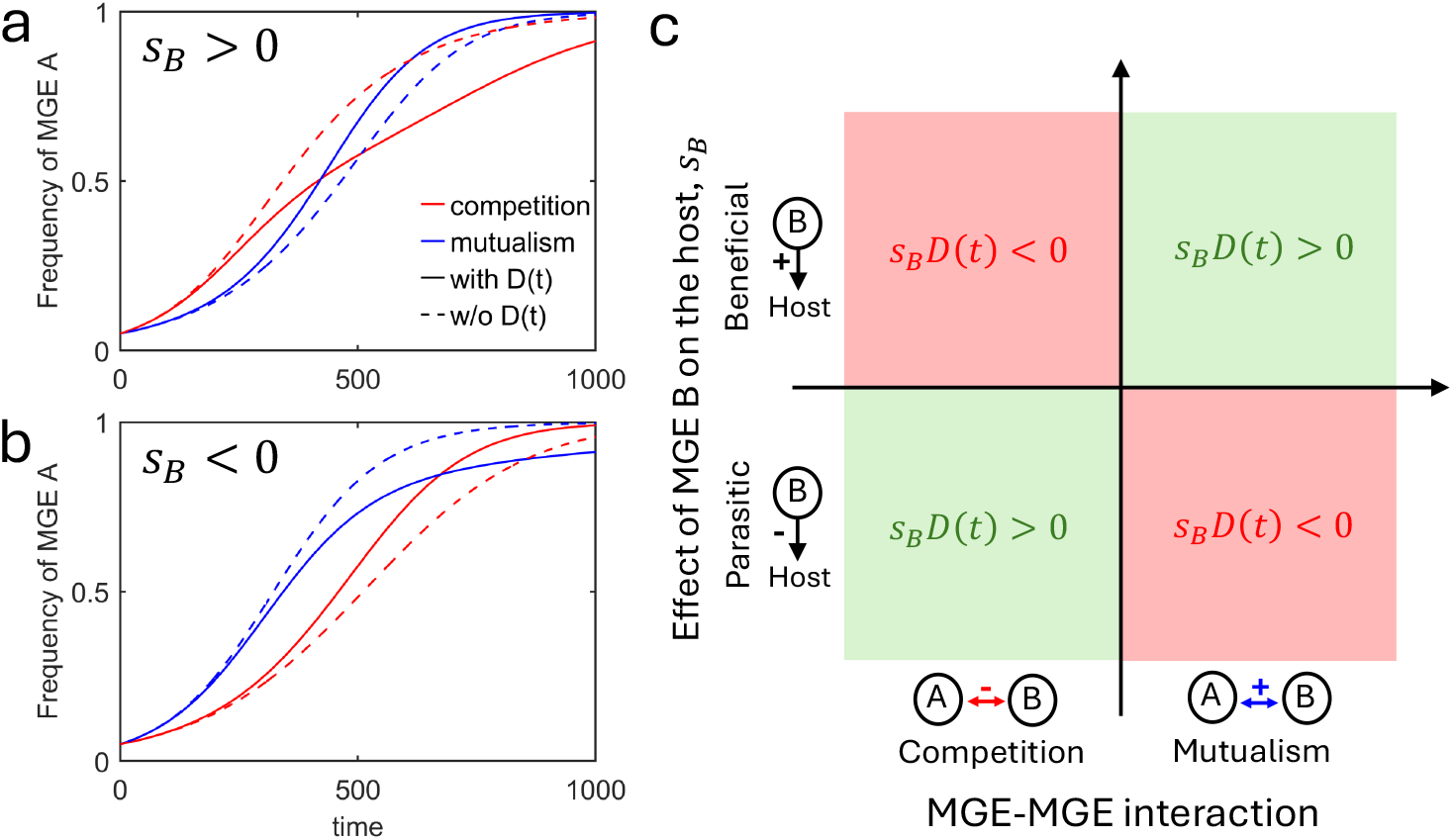
Influence of MGE-MGE interactions and MGE-host interactions on the speed of evolution of gene. *A*. Mutualistic MGE-MGE interactions generate positive associations between MGEs which increases the change in frequency of MGE *A* when MGE *B* is beneficial (panel **a**) and decrease it when MGE *B* is parasitic (panel **b**). Competitive MGE-MGE interactions, in contrast, generate negative associations between MGEs, which increase the change in frequency of MGE *A* when MGE *B* is parasitic panel **b**), and decrease it when MGE *B* is beneficial (panel **a**). The change in frequency of MGE *A* increases due to the indirect effect in the presence of MGE *B* when *s*_*B*_*D*(*t*) *>* 0 (green zone in panel **c**) and decreases in the presence of MGE *B* when *s*_*B*_*D*(*t*) *<* 0 (red zone in panel **c**). In panel **a**,**b**, all simulations assume MGE *A* is beneficial, *s*_*A*_ = 0.005, there is no SL, 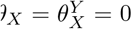, and we use the parameter values *b* = 4, *d* = 2, *β*_*AB*_ = 0, 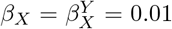 and initial conditions *p*_*A*_(0) = 0.05, *D*(0) = 0, *n*(0) = (*b* − *d*)*/b*. If HGT is competitive (red curves), 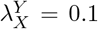 and *λ*_*X*_ = 0.9, whereas if HGT is mutualistic (blue curves), 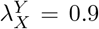 and *λ*_*X*_ = 0.1. Dashed lines indicate simulations when *D*(*t*) is forced to be equal zero. In panel **a**, *s*_*B*_ = 0.005 and *p*_*B*_(0) = 0.05, while in panel **b**, *s*_*B*_ = −0.005, and *p*_*B*_(0) = 0.95.

### 3.3 Long-term population response to environmental change

In the long-term, MGEs need not go to extinction or fixation, and can instead be stably maintained at an intermediate frequency. This can occur through opposing direct effects (e.g., the loss of MGEs through SL is counteracted by sufficiently strong HGT; Appendix A.1), or opposing direct and indirect effects (e.g., gene *Y* is beneficial and SL is mutualistic, or gene *Y* is parasitic and HGT is competitive; Appendix A.3). MGEs can also be maintained at an intermediate frequency through fluctuating or negative frequency dependent selection (NFDS) on MGE-linked genes. Indeed, there is evidence that a substantial portion of the accessory genome in some bacteria is under NFDS [26–28].

Importantly, the stable maintenance of MGEs at intermediate frequencies provides time for HGT and SL to build up *D*(*t*), creating an indirect effect. This indirect effect can then affect the evolutionary capacity of the bacteria population to respond to a change in the environment (e.g., an increase in antibiotics [5]; vaccination campaigns [26, 27]), affecting selection on one of the MGE-linked genes (e.g., gene *B* encodes for antibiotic resistance or is an antigen targeted by the vaccine). In what follows, we will assume the population has reached an equilibrium, then at time *t* = T_c_, there is a change in the environment affecting the direction and/or strength of selection on gene *B* by an amount Δ*s*_*B*_(*t*).

#### Interacting MGEs affect the short-term evolutionary response to environmental change

Immediately following the environmental change, the frequency dynamics of MGE *A* are

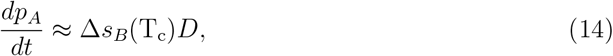

whereas MGE *B* will change in frequency according to Δ*s*_*B*_(T_c_)*p*_*B*_(*t*)*q*_*B*_(*t*). Thus the sign of *D*(*t*) dictates whether the change in selection on *B* has the same, or opposing (shortterm) consequences for the frequency of MGE *A* and *B*. In particular, if HGT and SL are competitive (*D*(*t*) *<* 0), then if MGE *A* increases (resp. decreases), MGE *B* decreases (resp. increases). Conversely, if HGT and SL are mutualistic, or the simultaneous gain and loss of MGEs is possible, if MGE *A* increases (resp. decreases), so will MGE *B*.

In the short-to-medium term, the microbial population most effectively responds to environmental change if *p*_*X*_(*t*) can change rapidly in the direction indicated by *s*_*X*_ (see equation (1)). From (6), this will occur if indirect selection on MGE *X, s*_*Y*_ *D*(*t*), shares sign with direct selection, *s*_*X*_. Thus, if selection is directional and constant, and Δ*s*_*B*_ and *s*_*A*_ share sign, mutualistic HGT and SL and/or the simultaneous gain and loss permit the microbial population to respond most rapidly to the environmental change, whereas if Δ*s*_*B*_ and *s*_*A*_ are of opposite sign, competitive HGT and SL permit the most rapid population response. The indirect effect of HGT and SL will remain the same as detailed in our previous analysis.

#### NFDS on MGE-linked genes slows microbial adaptation and reduces population resilience

Selection need not be constant nor directional. Therefore, suppose the genes are subject to NFDS of the form

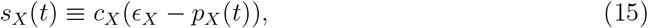

where *c*_*X*_ *>* 0 controls the relative strength of selection and *ϵ*_*X*_ *>* 0 is the optimal frequency of gene *X*. Phenomenological selection coefficients like equation (15) are a common assumption in genomic models of bacteria [5, 20, 26, 28]. Assume NFDS is strong relative to HGT and SL rates, and so *p*_*X*_(*t*) → *ϵ*_*X*_ for *t <* T_c_ (i.e., *s*_*X*_(T_c_) ≈ 0). Without loss of generality, suppose the environmental change increases the benefits of carrying gene *B*, Δ*s*_*B*_(T_c_) *>* 0 (so *ϵ*_*B*_ and potentially *c*_*B*_ increase). Then the indirect effect has two consequences (Fig. 6**c**). First, it will slow the change in frequency of MGE *B*, and second, it will perturb the dynamics of MGE *A*. This is because in the short-term, MGE *A* changes according to Δ*s*_*B*_*D*(*t*). This will mean *s*_*A*_(*t*) is no longer zero and instead has the opposite sign to *D*(*t*). As a result, indirect selection on MGE *B, s*_*A*_(*t*)*D*(*t*), will act counter to direct selection, Δ*s*_*B*_(*t*), slowing the change in frequency of MGE *B* (Fig. 6**b**). This also occurs for MGE *A*: direct selection, *s*_*A*_(*t*), and indirect selection, Δ*s*_*B*_(*t*)*D*(*t*), are of opposite sign, slowing the return of MGE *A* to equilibrium (Fig. 6**c**). Consequently, the time until the population recovers (i.e., returns to equilibrium), will increase, and thus HGT and SL will reduce what is sometimes referred to as the “resilience” of the population [25].

**Figure 6:**
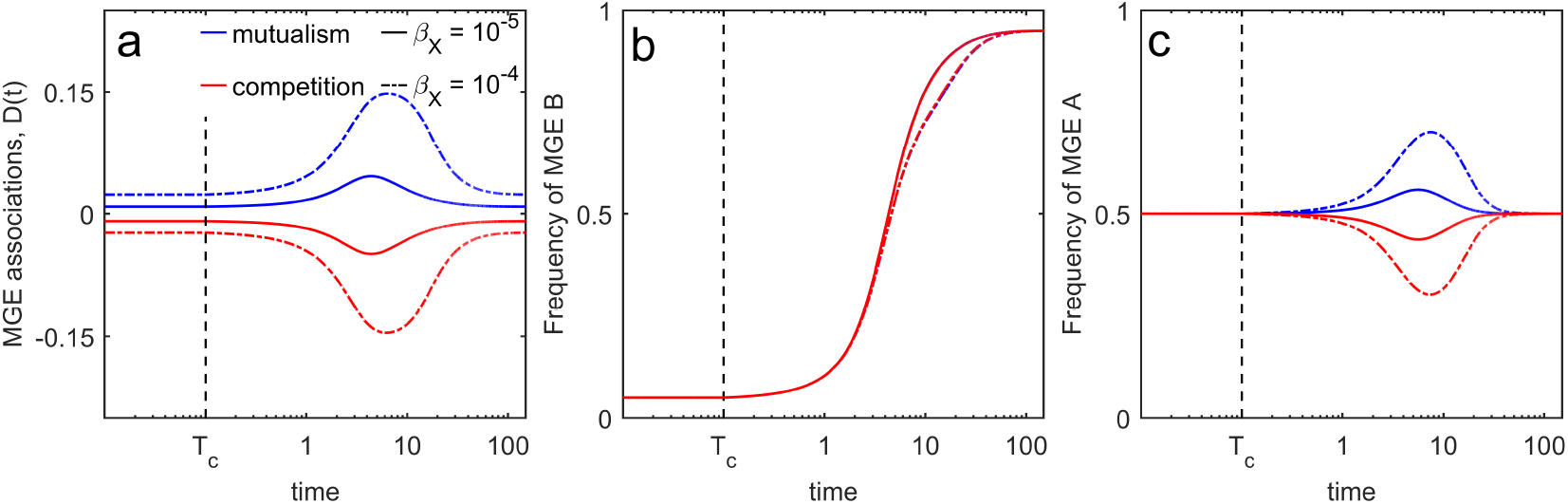
In the presence of NFDS, HGT and SL slow MGE evolution and reduce population resilience. In the long-term, HGT and SL can build up significant amounts of *D*(*t*) (panel **a**, *t <* T_c_). Following a change in environment at *t* = T_c_ (e.g., increase in antibiotics) affecting selection on gene *B*, the presence of *D*(*t*) will slow the change in frequency of MGE *B* (panel **b**) and reduce the population resilience, that is, increase the time required to return to the original state (panel **c**). In all simulations, *b* = 4, *d* = 2, *c*_*A*_ = *c*_*B*_ = 1, *ϵ*_*B*_ = 0.5, while for *t <* T_c_, *ϵ*_*A*_ = 0.05 and for *t >* T_c_, *ϵ*_*A*_ = 0.95. We then either assume *λ*_*X*_ = 0.05, 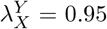 (solid curves), or *λ*_*X*_ = 0.95, 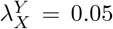 (dashed curves) while varying 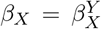 as indicated with *β*_*XY*_ = 0. Qualitatively similar predictions occur for SL.

## 4 Discussion

It is well understood that HGT and SL play an important role in microbial adaptation, speeding up or slowing down the change in frequency of MGE-linked genes as well as reshuffling genetic diversity across bacteria cells. What is less well understood is how interactions between MGEs affecting the likelihood, and rate, of HGT and SL shape microbial adaptation in the short- and long-term. Here, we develop a framework to disentangle these effects. We show MGE-MGE interactions affecting HGT and SL build up non-random associations between MGEs. If interactions are competitive (i.e., reduce HGT and increase SL), cells are more likely to carry a single MGE. If interactions are mutualistic (i.e., increase HGT and reduce SL) and/or the simultaneous gain and loss of MGEs is possible, cells tend to carry both MGEs or neither. These non-random associations affect the transient change in frequency of MGEs in two ways: by reducing the ability of HGT and SL to affect MGE frequency, and by producing indirect selection on MGE-linked genes. Owing to the combination of these effects, SL can speed the evolution of MGE-linked gene *X*, whereas HGT can slow its evolution, as compared to the situation when the gene is not MGE-linked (Appendix A.3).

In the long-term, HGT and SL can maintain MGEs at intermediate frequencies, even for neutral or deleterious MGEs [39, 40] and nonconjugative plasmids [41]. While the forces permitting the persistence of individual MGEs have been extensively investigated (thus resolving the so-called ‘plasmid paradox’ [11, 42–45]), less attention has been paid to how MGE-MGE interactions affect MGE persistence [19]. Our framework clarifies that in order for MGEs to persist due to MGE interactions, the indirect effect, mediated through non-random MGE associations, must oppose the direct effect. For example, if gene *B* is beneficial and SL is mutualistic, or gene *B* is parasitic and HGT is competitive, MGE interactions can maintain MGEs at intermediate frequencies (Appendix A.3). The long-term presence of non-random MGE associations affects the ability of microbial populations to respond to environmental change. If the MGE-linked genes are under NFDS, MGE-MGE interactions affecting HGT and SL reduce the speed of microbial adaptation and reduce population resilience [25].

### MGE-MGE interactions and epistasis

To isolate the role of MGE-MGE interactions affecting HGT and SL, we ignored the possibility of epistasis between genes affecting per-capita growth rate. However, epistasis in per-capita growth rate is empirically [2, 15, 46–48] and theoretically [49, 50] observed in MGE-bacteria dynamics. Previous work has shown that epistasis can both slow [8], and speed [51] the evolution of antibiotic resistance, as well as produce strain-specific variability in MGE (plasmid) evolution [10]. More broadly, epistasis in per-capita growth rate is well-known to be an important contributor to the generation and build-up of non-random associations between genes [52] and so can function similarly to MGE-MGE interactions affecting HGT and SL. Indeed, when HGT and SL do build-up *D*(*t*), this can be thought of as epistatic interactions in HGT and SL rates. Although MGE-MGE interactions affecting HGT and SL can generate and maintain non-zero *D*(*t*), and thus mimic the action of epistasis in per-capita growth rate, such interactions do not directly affect the per-capita growth rate of the bacterial population (equation (1)). Consequently, MGE-MGE interactions in HGT and SL are not equivalent to epistasis in per-capita growth rate.

That different types of interactions build-up *D*(*t*) presents a challenge to interpreting the source of pervasive non-random MGE associations. For example, an analysis by San Millan et al. [47] found an over-representation of cells carrying multiple plasmids across different phyla (*D*(*t*) *>* 0). While experiments in *Pseudomonas aeruginosa* suggest this may be due to positive epistasis minimizing the fitness costs associated with carrying multiple plasmids [47], it is not known whether this holds true more broadly. Our analysis suggests an alternative possibility is MGE interactions affecting HGT and SL: if plasmids can be simultaneously lost or gained, this will produce an over-representation of cells carrying multiple plasmids unless HGT and SL are strongly competitive. Thus pervasive positive epistasis in fitness costs is not necessary to explain the distribution of plasmids across phyla [47].

### NFDS, HGT and the dynamics of accessory genomes

Empirical evidence suggests a significant portion of the accessory genome of some bacteria species, including *Streptococcus pneumoniae*, is under stabilizing selection [28], and considerable effort has been paid to understand how this effects microbial evolution [20, 26–28]. Harrow et al. [20] argue the combination of NFDS and the possibility the loss of accessory genes during HGT is more probable than the gain (‘asymmetric transformation’ [20]), could explain bacteria accessory genome patterns, particularly the differentiation into distinct strains. In our model, asymmetric transformation corresponds to the possibility of simultaneous transmission of multiple MGEs, with the probability of establishment independent of the identity of the donor. This will produce positive linkage disequilibrium between accessory genes. As this will lead to an over-representation of certain genic combinations, such a process will generate ‘strains’ [53], but our analysis here predicts these strains correspond to cells carrying either numerous accessory genes or very few, producing a bimodal distribution of the ‘length’ of accessory genomes, a testable prediction.

### HGT and SL and the population response to vaccination

That MGE-MGE interactions reduce population resilience has implications for predicting the response of bacteria populations to serotype-specific vaccination campaigns [26–28]. These models consider pathogen genomes consisting of a number of loci, one of which is the serotype, while the others are accessory genes under NFDS at an ‘optimum’ (*s*_*X*_ ≈ 0) corresponding to their observed frequency prior to vaccination. Serotype-specific vaccination then removes all genomes from the population belonging to vaccine serotypes, perturbing the accessory genes from their optimum. These studies cleverly reason that the subsequent rebound of the population post-vaccination is driven by the return of the NFDS genes to their prevaccine ‘optima’, with the serotype frequencies changing due to genetic hitchhiking [26, 27]. Our analysis reveals two challenges to this approach. First, while such methods necessarily account for non-random associations between serotype and the individual accessory genes, they do not account for non-random associations between the accessory genes, due to, for example, MGE-MGE interactions. The presence of non-random accessory gene associations will mean gene frequencies are insufficient to predict the resulting dynamics. Second, the presence of *D*(*t*) from MGE-MGE interactions before the environmental change means the population need not return to its previous state over biologically relevant timescales, as indirect selection will both perturb allele frequencies, and slow their speed of evolution (return to original state).

### Incompatibility groups and what genes are carried by plasmids

Incompatibilities between plasmids are often modeled as a reduced probability of establishment 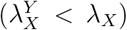 with no co-transmission (*β*_*AB*_ = 0), they could equally be modeled as increasing rates of SL 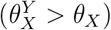. If, in addition, the simultaneous loss is possible, *θ*_*AB*_ *>* 0, whether plasmids are less (*D*(*t*) *<* 0), or more (*D*(*t*) *>* 0), likely to be found in the same cell than expected will depend not only on the frequency of the MGEs, but the strength of the incompatibility. This stands in contrast to a situation in which plasmids are unable to co-infect the same cell, which always produces negative *D*(*t*).

This has relevance for theoretical work seeking to understand why certain genes (e.g., antibiotic resistance [3] or cooperative traits [54–58]) are more likely to be carried by an MGE [4, 6, 29, 59, 60]. Models of this type usually assume there are two competing plasmids, one that carries the (beneficial) gene of interest (say MGE *A*) and one that does not (MGE *B*). As the plasmids are assumed to differ only at the gene of interest, they are taken to belong to incompatibility groups [33], and this is modeled by assuming HGT is blocked whenever the other plasmid is present in the recipient (i.e., 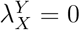, *β*_*AB*_ = 0). Assuming plasmid *B* is parasitic (as it does not carry the beneficial allele), our analysis indicates this type of HGT can speed the evolution of gene *A* through indirect selection. However, the conclusions may be reversed if incompatibilities affect rates of SL.

#### HGT and SL and the community response to antibiotic treatment

Recent work suggests HGT increases the stability of diverse bacterial communities against antibiotic treatment [5]. The framework developed here can be modified to provide insight into how dependencies in HGT and SL affect the community response to antibiotic treatment. In particular, suppose we have two species under stabilizing ecological selection and that antibiotic resistance is MGE-linked and is stably present at some low frequency in both species (so *s*_*X*_(*t*) are of the form (15)). As HGT rates often show a positive correlation with genetic similarity between donor and recipient [24, 35, 61–63], suppose the rate of within-species HGT is greater than the rate of between-species HGT. Under these circumstances, antibiotic resistance is more likely to be carried by the more abundant species, and this likelihood increases with genetic distance between species (Appendix A.4).

Our analysis suggests this will have two consequences. First, increasing the genetic diversity of a bacterial community will increase the amount of *D*(*t*), slowing antibiotic resistance evolution. Second, this will also slow the recovery of the community to its original species composition. Thus after accounting for genetic differences between species, HGT can decrease the stability and resilience of complex communities. The role of biases in HGT rates represent an important consideration for future work, in addition to considering the role of species interactions (e.g., competition, cooperation, etc.) as done by Coyte et al. [5]. The framework established here is well-suited to unravel these dynamics, and can be extended to include multiple MGEs in a multi-species bacteria community.

## Conclusion

In classical population genetics, recombination acts to shuffle alleles, thereby breaking up non-random allelic associations. As our framework reveals, this is generally not true for the action of HGT and SL: both HGT and SL are likely to generate nonrandom associations between MGEs. By affecting the distribution of MGEs in the population, HGT and SL can either speed up or slow down the change in frequency of MGE-linked genes. More generally, the framework developed here provides a useful starting point for understanding how genetically diverse bacteria populations are likely to respond to environmental changes, such as antibiotic treatment or vaccination campaigns.

## Acknowledgements

We thank Eduardo Rocha for his many useful comments on an earlier version of the manuscript.

## Box 1: Fitness interactions involving MGEs

Each MGE may affect the fitness of its host as well as the fitness of other MGEs. In the following, we classify the different interactions based on their fitness consequences for the parties involved. We illustrate each case with an example.

**MGE-Host interactions**

A MGE carrying gene *X* may affect the per capita rate of cell division or death of its host by an amount *s*_*X*_. The sign of *s*_*X*_ distinguishes between two types of MGEs:

- The MGE is *parasitic* if it is costly for its host (*s*_*X*_ *<* 0). For example, many bacteriophages decrease the survival and reproduction of an infected host.
- The MGE is *beneficial* if it is advantageous for its host (*s*_*X*_ *>* 0). For example, a plasmid carrying an antibiotic resistance gene increases the survival of its host in the presence of the antibiotic.

**MGE-MGE interactions**

A MGE (say *Y* ) may also interfere with the spread and loss of another MGE (say *X*) during transmission (if 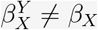), establishment (if 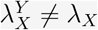) and/or segregation loss (if 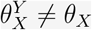). Such MGE-MGE interactions are of two main types:

- MGE-MGE interactions are *competitive* if the presence of the other MGE decreases transmission 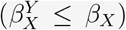, decreases establishment 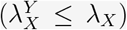, and/or increases loss 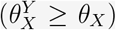. For example, the presence of plasmid *Y* may reduce the ability of plasmid *X* to colonize the host cell, 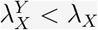.
- MGE-MGE interactions are *mutualistic* if the presence of the other MGE increases transmission (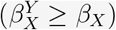), increases establishment 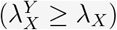, and/or decreases loss 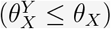. For example, some plasmids benefit from co-transmission.

Both competitive and mutualistic interactions may be asymmetric (i.e., in the extreme, amensalism or commensalism). For example, an (asymmetric) mutualism occurs if one MGE (the hitcher) relies on the presence of a helper MGE for transmission, but the helper MGE is only weakly affected by the hitcher.

## A Appendix

### A.1 Model

Here we provide the equations describing the dynamics of the densities of the different bacteria cell types. To do so, we first detail the assumptions that lead to the specification of per-capita growth, segregation loss, and horizontal gene transfer.

#### Per-capita growth

Let *n*_*ij*_(L,*t*) denote the density of bacteria cells of type *ij* ∈ {∅∅, *A*∅, ∅*B, AB*} at time *t* and 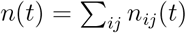 denote the total density of bacteria cells, where *X* = {*A, B*} indicates the presence of gene *X* (and so carriage of MGE *X*), while a ∅ indicates the absence of the corresponding MGE. Bacteria cells divide at a density-dependent per-capita rate *b*(1−*n*(*t*)), and die at a per-capita rate *d*. In addition, carriage of MGE *X* affects the rate of cell division and bacteria cell death by an amount *s*_*X*_(*t*). If *s*_*X*_(*t*) *>* 0, carriage of MGE *X* increases the bacterial per-capita growth rate, and so MGE *X* is *beneficial* for the bacteria, whereas if *s*_*X*_(*t*) *<* 0, carriage of MGE *X* decreases the bacterial per-capita growth rate and so MGE *X* is *parasitic*. We leave *s*_*X*_(*t*) general for the moment, and so *s*_*X*_(*t*) may fluctuate with time (e.g., due to changes in antibiotic concentration), but may also depend on bacteria population densities if, for example, carriage of MGE *X* affects the rate of cell division. We can write the per-capita growth rate of bacteria cell type *ij* as

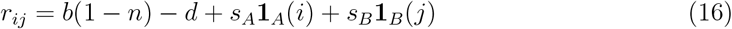

where **1**_*X*_(*k*) is an indicator variable, equal to 1 if *X* = *k* and 0 otherwise.

##### Segregation loss

During cell division bacteria cells may lose MGEs due to segregation loss (SL). The probability of SL may depend on whether the other MGE is present or not, and if present, whether both MGEs are simultaneously lost from the cell. To capture these possibilities we let *θ*_*X*_ denote the probability that MGE *X* is lost from a cell that does not carry the other MGE *Y* , and 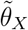 denote the probability that only MGE *X* is lost from a cell that does carry the other MGE *Y* . Let *θ*_*AB*_ denote the probability that a cell carrying both MGEs loses them simultaneously. Thus, the probability that a cell carrying both MGEs loses MGE *X* per cell division is 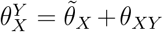 , where 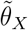 refers to the probability that a cell carrying both MGEs loses only MGE *X*, while the probability that a cell carrying both MGEs loses at least one MGE per cell division is 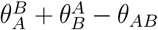. SL produces five possible changes to bacteria cells:

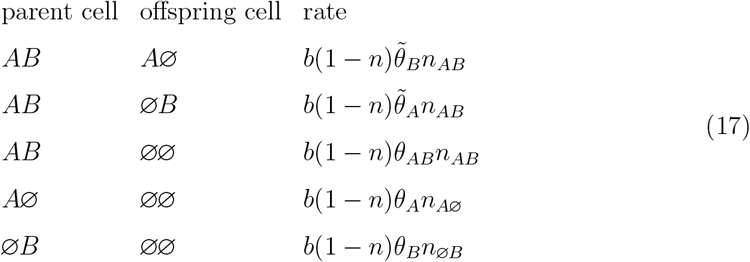

Using these transitions, the net change in bacterial cell type *ij* ∈{∅∅, *A*∅, ∅*B, AB*} due to SL is

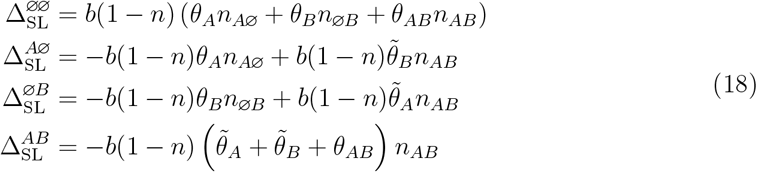

We define the probability of SL per cell division of a randomly selected cell carrying MGE *A* as

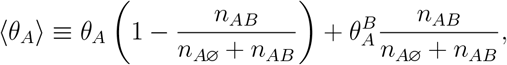

while for MGE *B*, we have

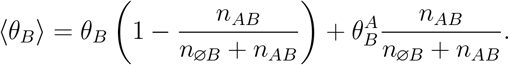

##### Horizontal gene transfer

MGEs may be transferred from one bacteria cell (henceforth, the “donor”) to another (henceforth, the “recipient”) through horizontal gene transfer (HGT). We assume that HGT can be decomposed into two components: (1) the transmission of the MGE(s) from the donor, and (2) the successful establishment of the MGE(s) in the recipient.

Transmission of the MGE(s) may depend on whether the donor carries multiple MGEs, and if so, whether one or both are transmitted. Let *β*_*X*_ denote the transmission rate of MGE *X* if the donor does not carry the other MGE *Y* , and 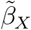 denote the transmission rate of only MGE *X* if the donor carries the other MGE *Y* . Let *β*_*AB*_ denote the rate of transmission of both MGEs *A* and *B*. Thus the total rate at which MGE *X* is transmitted from a host that carries both MGEs is 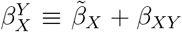 . The total rate at which a host carrying both MGEs transmits at least one is 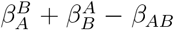, while the average rate of transmission of MGE *A* by a randomly selected cell carrying MGE *A* is *n*⟨*β*_*A*_⟩, where

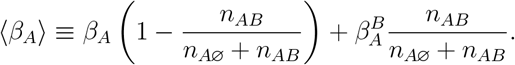

Similarly, the average rate of transmission of MGE *B* by a randomly selected cell carrying MGE *B* is *n*⟨*β*_*B*_⟩, where

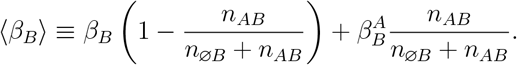

The successful establishment of the MGE(s) in the recipient may depend on the transmitted MGE(s), as well as the MGEs the donor carries. Let 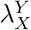 denote the probability of successful establishment of MGE *X* when the recipient carries the other MGE *Y* , and let *λ*_*X*_ denote the probability of establishment if the donor does not. If both the MGEs are transmitted, we assume that whether one establishes or not does not affect the establishment of the other. Thus, both transmitted MGEs establish with probability *λ*_*A*_*λ*_*B*_. For simplicity, we ignore “dose” or “multicopy” effects, and so if a MGE is transmitted to a host who already carries that MGE, the state of the recipient host is unchanged, irrespective of whether establishment is successful or not. The probability of successful establishment of MGE *A* in a randomly selected host that does not carry MGE *A* is

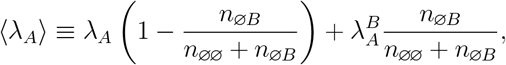

while the probability of successful establishment of MGE *B* in a randomly selected host that does not carry MGE *B* is

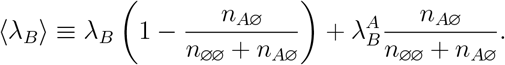

HGT thus produces nine possible changes to bacteria cells:

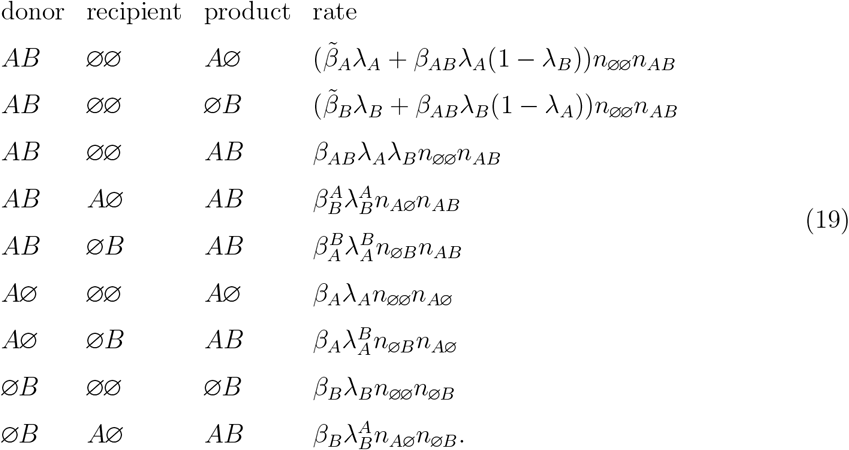

Using these transitions, the net change in bacterial cell type *ij* ∈{∅∅, *A*∅, ∅*B, AB}* due to HGT is

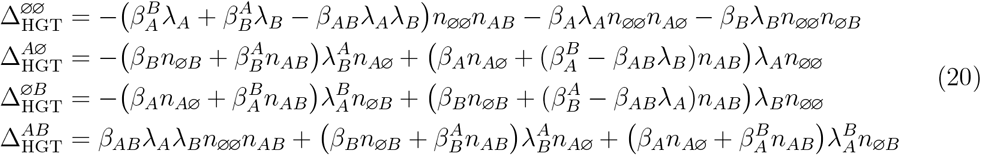

##### Equations describing the dynamics of the different bacteria cell types

In combination, the dynamics of the density of bacteria cell type *ij* is

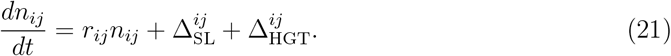

The dynamics of the total density of bacteria cells is:

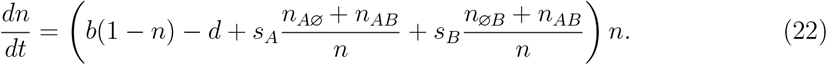

##### Equations describing the evolutionary dynamics

The frequency of MGE *A* and *B* are defined as

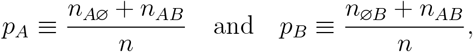

while the non-random associations between MGES is

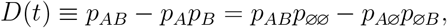

where *p*_*ij*_ = *n*_*ij*_*/n*. The dynamics of the frequency of MGE *X* are

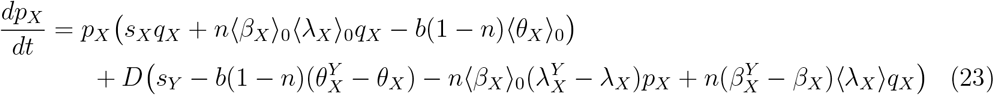

where *q*_*X*_ = 1 −*p*_*X*_ and ⟨ *Z*_*X*_⟩ _0_ for *Z* ∈ {*β, λ, θ}* means we evaluate the quantity ⟨ *Z*_*X*_⟩when *D*(*t*) = 0. From (23), notice that in the absence of selection the balance between the direct effects of HGT and SL can maintain the MGEs at an intermediate frequency if the inequality

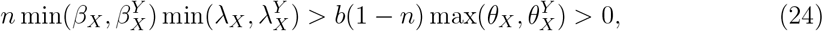

is satisfied, that i s, the rate of HGT exceeds the rate of SL when the MGE is rare.

The full equation for dD/dt is

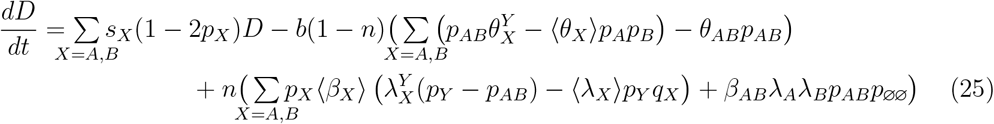

#### A.2 Generation of MGE associations, *D*(*t*)

First, consider how HGT and SL affect the generation of MGE associations, *D*(*t*). From equation (25) if we set *D*(*t*) = 0, we obtain

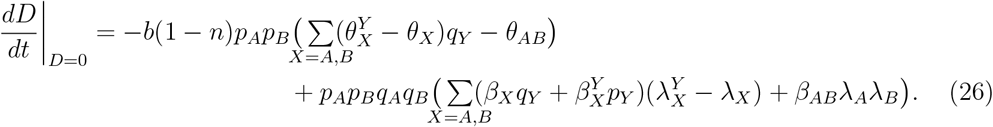

##### Segregation loss

From equation (26), SL generates positive *D*(*t*) if

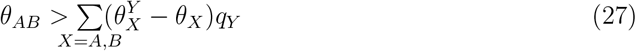

and negative *D*(*t*) if the inequality is reversed.

In the main text, we observe that if *θ*_*AB*_ *>* 0 and 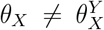 , then the epidemiology of the MGEs plays an important role in determining whether the dependencies of SL on genetic background (difference between *θ*_*X*_ and 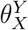 ) matter or not for the generation of *D*(*t*). In particular, if the MGEs are common, *q*_*Y*_ (*t*) → 0, then SL dependencies on genetic background do not matter, whereas if MGEs are rare, *q*_*Y*_ (*t*) → 1, SL dependencies on genetic background do matter. In the main text, we argue this is because when MGEs are very common (*q*_*Y*_ (*t*) → 0), the only SL event that is both frequent and has a non-negligible impact on *D*(*t*) is when *AB* cells lose both MGEs. Thus all that matters is that *θ*_*AB*_ *>* 0. On the other hand, when MGEs are very rare (*q*_*Y*_ (*t*) → 1), although *A*∅ and ∅*B* cells are more frequent and so more likely to undergo SL events than *AB* cells, *AB* cells undergoing SL have a much larger impact on *D*(*t*). Consequently, all SL transitions must be taken into account for the generation of *D*(*t*), and so SL dependencies on genetic background matter. To show this, first consider the case in which the MGEs are very common, *q*_*Y*_ (*t*) → 0.

In this circumstance, most of the population are *AB* cells, and so at linkage equilibrium (*D*(*t*) = 0), we have *p*_*AB*_(*t*) ≈1, *p*_*A*∅_(*t*) ≈*p*_∅*B*_(*t*) = *ω* and *p*_∅∅_ ≈*ω*^2^, where 0 *< ω* ≪1. Since *ω* is very small, virtually all SL events involve *AB* cells and so are our focus. *AB* cells can undergo three possible transitions, simultaneous loss of both MGEs, or loss of either MGE, but not the other. Suppose over a small increment of time ∆*t*, a fraction ∆*p* of *AB* cells lose both MGEs. For such transitions, we can write the change in *D*(*t*) as

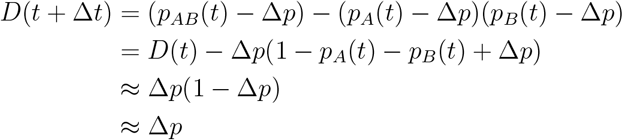

after we have let *p*_*X*_(*t*) ≈ 1 and noted that the population is initially in linkage equilibrium, *D*(*t*) = 0. For *AB* → *A*∅ transitions, we can write the change in *D*(*t*) as

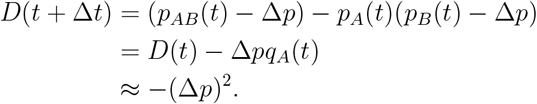

Thus when *q*_*X*_(*t*) ≈ 0 (i.e., *q*_*X*_(*t*) ≈ ∆*p*), *AB* → *A*∅ and *AB* → ∅*B* transitions have a negligible effect on *D*(*t*) relative to *AB* → ∅∅ transitions. Consequently, if MGEs are very common and *θ*_*AB*_ *>* 0, SL will generate positive *D*(*t*).

If MGEs are rare, *q*_*Y*_ (*t*) → 1, than most of the population are ∅∅ cells. Thus at linkage equilibrium, *p*_∅∅_(*t*) ≈1, *p*_*A*∅_(*t*) = *p*_∅*B*_(*t*) ≈*ω*, and *p*_*AB*_(*t*) ≈ *ω*^2^ for 0 *< ω*≪ 1. This implies that the rate of SL events involving cells carrying a single MGE is an order of magnitude higher than SL events involving cells carrying two MGEs (*ω* vs *ω*^2^). However, consider the change in *D*(*t*) over a small amount of time ∆*t*, and the subsequent rate of the event:

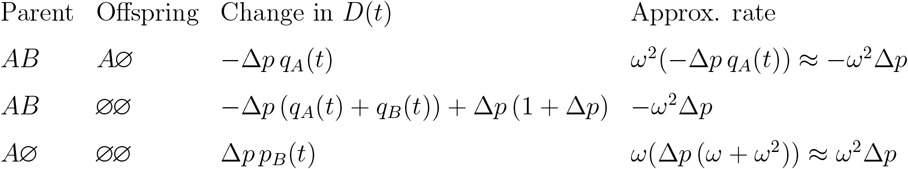

Consequently, to leading order all the approximate rates are of comparable order of magnitude, *ω*^2^∆*p*. Hence, although *A*∅ and ∅*B* cells are much more abundant than *AB* cells (*ω* vs *ω*^2^), because SL events involving *AB* have a significantly larger impact on *D*(*t*) when MGEs are rare, the generation of *D*(*t*) will be affected by the relative values of all the parameters, 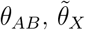 , and *θ*_*X*_.

##### A.2.1 Horizontal gene transfer

From equation (26), HGT generates positive *D*(*t*) if

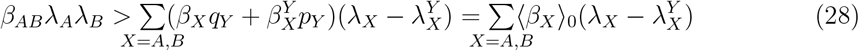

and negative *D*(*t*) if the inequality is reversed.

#### A.3 Indirect selection

Next, we consider in more detail the role of indirect selection. To do so, we separately consider SL and HGT, and we suppose there is no direct selection on MGE *A*.

##### Segregation loss and indirect selection

First, consider the interaction between SL and indirect selection. Here we have

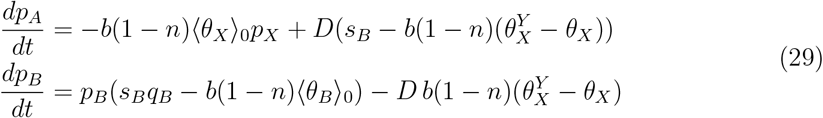

First, note that the necessary condition for MGE *B* to increase when rare is

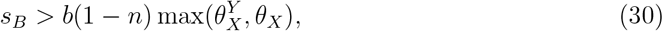

where *n* = (*b* − *d*)*/b*. If MGE *B* goes extinct MGE *A* will as well. It need not be true, however, that if MGE *B* avoids extinction so will MGE *A*.

Suppose that the simultaneous loss of MGEs is not possible. Then the sign of *D*(*t*) is opposite to the sign of 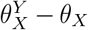. Consequently, indirect selection will speed the loss of MGE *A* if *s*_*B*_ is of opposite sign to 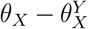 , and so MGE *A* goes extinct. If *s*_*B*_ is of the same sign as 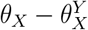 , indirect selection slows the loss of MGE *A*. In this case, if gene *B* is sufficiently beneficial such that MGE *B* will avoid extinction, it is possible that indirect selection will be sufficiently strong that MGE *A* can avoid extinction and stably persist in the population.

If the simultaneous loss of MGEs is possible, then if gene *B* is beneficial (resp. deleterious), indirect selection slows (resp. speeds) the loss of MGE *A*. Notably, even if gene *B* is beneficial, long-term persistence of MGE *A* is not possible. This is because if 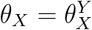 , then ⟨*θ*_*X*_ ⟩_0_ = *θ*_*X*_. Consequently, the dynamics of *p*_*B*_(*t*) and *n*(*t*) do not depend on either *p*_*A*_(*t*) or *D*(*t*), and so will ultimately go to an equilibrium unaffected by the presence or dynamics of MGE *A*. The equilibrium at which MGE *B* persists is:

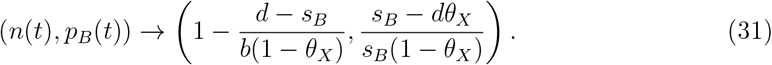

It is necessary that *d > s*_*B*_ (because MGE *B* reduces death rate), while in order for equilibrium (31) to be valid, *s*_*B*_ *> dθ*_*X*_. At the equilibrium (31), the ODEs for *p*_*A*_(*t*) and *D*(*t*) are linear in *p*_*A*_(*t*) and *D*(*t*), and so if we compute the Jacobian for *dp*_*A*_*/dt, dD/dt* evaluated at equilibrium (31), which we denote *J*_*SL*_, it can be shown that

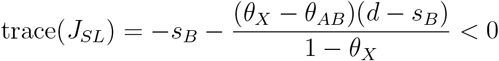

and

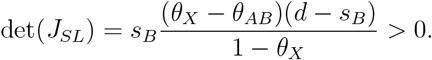

Therefore, ultimately MGE *A* will go extinct.

##### Horizontal gene transfer and indirect selection

Next, consider the interaction between horizontal gene transfer and indirect selection. If we focus upon the forces capable of generating *D*(*t*) (so set 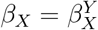 ), we obtain

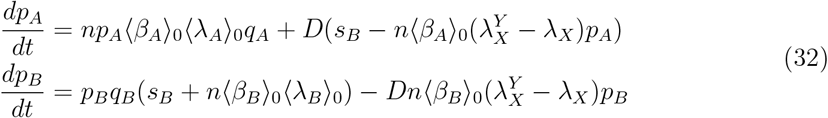

The sufficient condition for MGE *B* to not go to fixation in the long-term is that

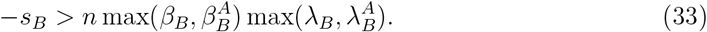

Suppose first that the simultaneous gain of MGEs is not possible, *β*_*AB*_ = 0. Since 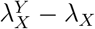 shares sign with *D*(*t*), if *s*_*B*_ shares sign with (resp. is of opposite sign to) 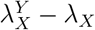, indirect selection increases (resp. decreases) the frequency of MGE *A*. If indirect selection decreases the frequency of MGE *A*, and in addition, MGE *B* is sufficiently parasitic such that it is present at an intermediate frequency, indirect selection can cause MGE *A* to stably persist at an intermediate frequency.

If the simultaneous gain of MGEs is possible, then if gene *B* is beneficial (resp. deleterious), indirect selection speeds (resp. slows) the fixation of gene *A*. As was the case for the simultaneous loss of MGEs, if MGEs are simultaneously gained, MGE *A* (and so gene *A*) cannot persist at an intermediate frequency and instead go to fixation. To show this, note that 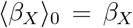 and 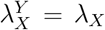 so the dynamics of *p*_*B*_(*t*) and *n*(*t*) will not depend upon *D*(*t*) and *p*_*A*_(*t*). Suppose MGE *B* persists at an intermediate frequency (so inequality (33) is satisfied), in which case the equilibrium value of *p*_*B*_ satisfies *s*_*B*_ = −*np*_*B*_*λ*_*X*_*β*_*X*_ from setting *dp*_*B*_*/dt* = 0. Next, we compute the Jacobian for *dp*_*A*_*/dt, dD/dt* evaluated at the equilibrium for which *B* is present at an intermediate value and (*p*_*A*_, *D*) = (1, 0). Denote this matrix *J*_*HGT*_ . Then we have

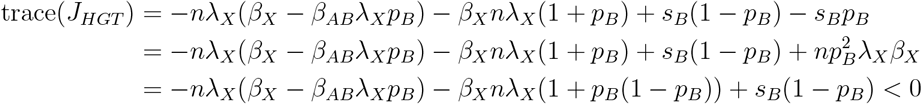

where we used that *s*_*B*_ = −*np*_*B*_*λ*_*X*_*β*_*X*_ and that *β*_*X*_ ≥ *β*_*AB*_ ≥ *β*_*AB*_*λ*_*X*_*p*_*B*_. Likewise,

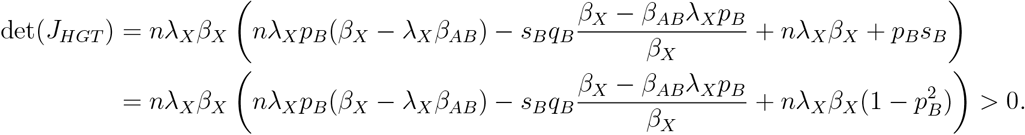

So this equilibrium is asymptotically stable, and hence gene *A* goes to fixation.

##### Indirect selection and the speed of evolution

If we consider the speed of evolution of a beneficial gene *A*, the indirect effect can mean that the net consequence of HGT is to slow its evolution, whereas SL can speed its evolution, as compared to if *A* is not MGE-linked. To see this, focus upon competitive or mutualistic SL and HGT. If 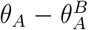 and *s*_*B*_ share sign, the indirect effect of SL on MGE *A* and indirect selection on MGE *B* work together to increase the frequency of MGE *A*. This can cause genes undergoing SL to evolve faster (Fig. A.1). If 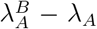 is of opposite sign to *s*_*B*_, the indirect effect of HGT on MGE *A* and indirect selection on MGE *B* work together to decrease the frequency of MGE *A*. This can cause MGE-linked genes undergoing HGT to evolve slower (Fig. A.1). If instead MGEs can be simultaneously gained or lost, the indirect effect is limited to indirect selection, and so it is less likely that HGT can slow or SL can speed evolution.

#### A.4 Alternate models of HGT

In the main text we briefly discuss an example in which one locus, {*B*_1_, *B*_2_} indicates species identity, while the other locus, {∅, *A*} indicates whether or not a cell is infected by a plasmid that carries antibiotic resistance. Letting 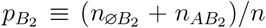 denote the frequency of species 2, we suppose that the species are maintained in the population due to stabilizing ecological selection,

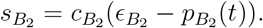

**Figure A.1:**
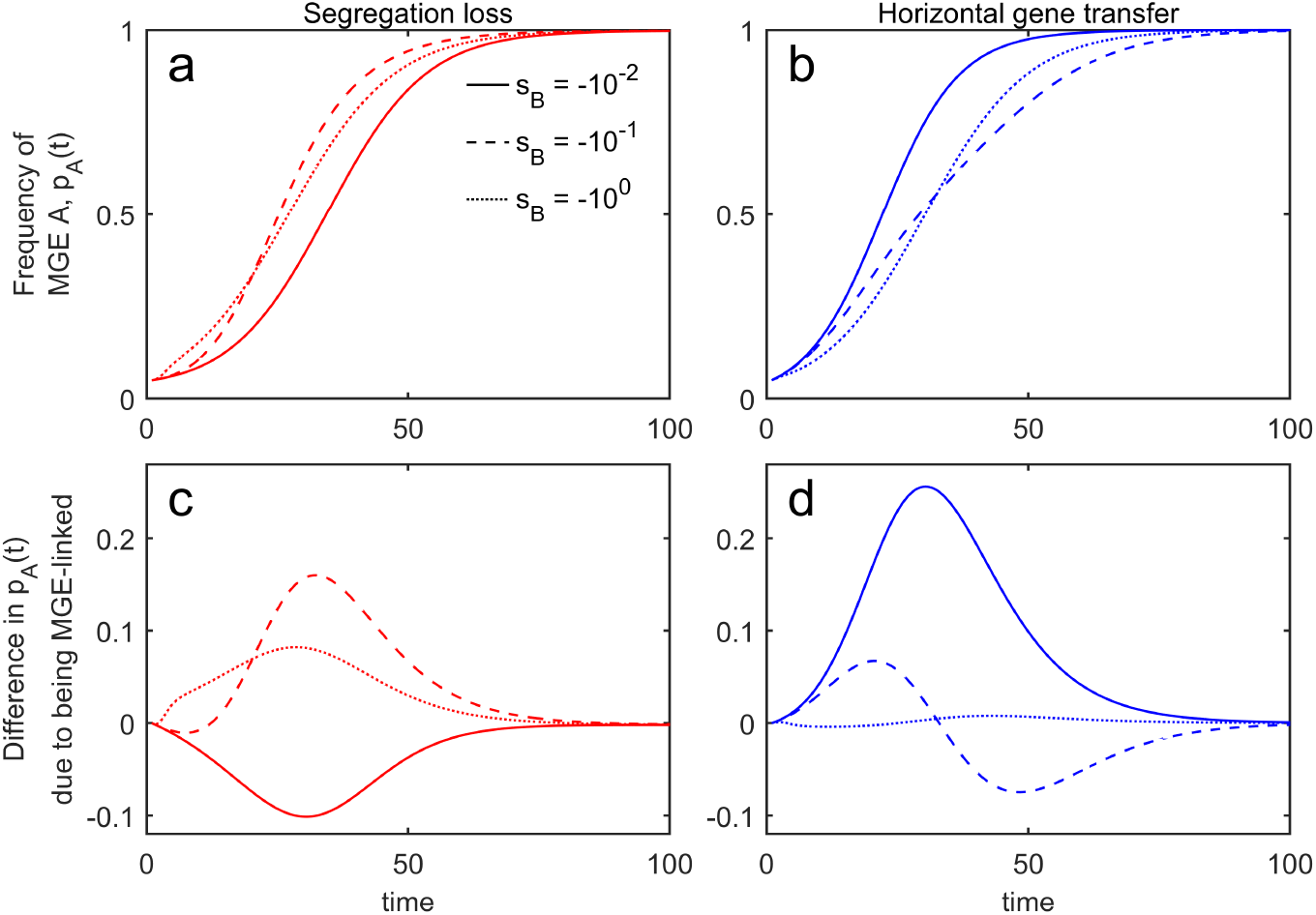
SL and HGT can both speed and slow the evolution of genes under directional selection. The generation of *D*(*t*) by competitive SL and mutualistic HGT can reverse the expectation that SL slows down the spread of MGEs (panels **a**,**c**) and HGT speeds up the spread of MGEs (panels **b**,**d**). In all panels we fix the strength of selection on MGE *A* and vary the degree to which MGE *B* is parasitic. In all simulations, *b* = 4, *d* = 2, *s*_*A*_ = 10^*−*1^, and *p*_*A*_(0) = 0.05, *p*_*B*_(0) = 0.85, *D*(0) = 0, *n*(0) = (*b* + *s*_*B*_*p*_*B*_(0) − *d*)*/b*. In panels **a**,**c**, 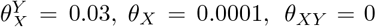, and 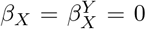. In panels **b**,**d**, 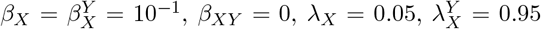and 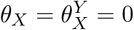.

This could arise if, for example, the species dynamics were modeled using Lotka-Volterra equations. In this example, we allowed for the possibility HGT rates may depend upon genetic similarity between the donor and the recipient. In particular, if the donor is the same species as the recipient, HGT of MGE *A* occurs at rate *β*_*ws*_ (*ws* for within-species), whereas if the donor is not the same species as the recipient, HGT of MGE *A* occurs at rate *β*_*bs*_ (*bs* for between-species). As genetic similarity between cells tends to increase HGT rates, we assume *β*_*ws*_ *< β*_*bs*_. This gives rise to the following four transitions through HGT:

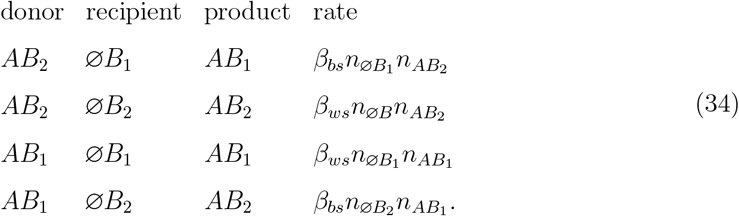

Next, we define

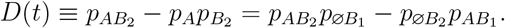

Thus *D*(*t*) captures whether antibiotic resistance is over-represented in species *B*_2_ (*D*(*t*) *>* 0), or species *B*_1_ (*D*(*t*) *<* 0). The equation for the change in *D*(*t*) can be written

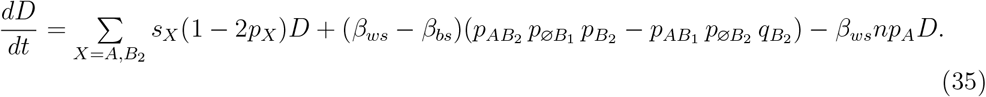

At linkage equilibrium, *D*(*t*) = 0, this yields

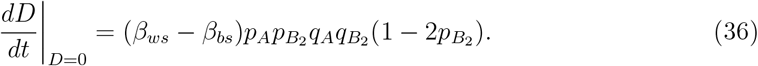

Since *β*_*ws*_ *> β*_*bs*_, if 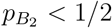, HGT will generate negative *D*(*t*) (species *B*_1_ is more likely to carry antibiotic resistance), whereas if 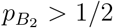, HGT will generate positive *D*(*t*) (species *B*_2_ is more likely to carry antibiotic resistance). Consequently, the more abundant species is more likely to carry antibiotic resistance.

